# Lrig3 restricts the size of the colon stem cell compartment

**DOI:** 10.1101/2022.03.08.483523

**Authors:** Janelle G. Stevenson, Ryan Sayegh, Natalie Pedicino, Natalie A. Pellitier, Tim Wheeler, Matthew E. Bechard, Won Jae Huh, Robert J. Coffey, Anne E. Zemper

**Author notes:** Corresponding Author: Anne E. Zemper, PhD, 1229 University of Oregon, 1318 Franklin Avenue, Room 273 Onyx Bridge, Eugene, OR 97403.

## Abstract

The cellular census of the colonic crypt is tightly regulated, yet the molecular mechanisms that regulate this census are not fully understood. Lrig3, a transmembrane protein, is expressed in colonic crypt epithelial cells, including the stem, progenitor, and differentiated cell types. Mice missing Lrig3 have a disruption in their cellular census: using a novel *Lrig3*^*-/-*^ mouse we demonstrate that *Lrig3*^*-/-*^ mice have more cells per crypt, a greater mucosal area, and longer colons compared to wildtype mice, suggesting the expression of Lrig3 is required for both the total number of epithelial cells in the mouse colon, as well as colon length. In addition, we show *Lrig3*^*-/-*^ mice have significantly more stem, progenitor, and deep crypt secretory cells, yet harbor a normal complement of enteroendocrine, Tuft, and absorptive cells. *Lrig3*^*-/-*^ mice also have a concomitant decrease in phosphorylated Extracellular signal-related kinases, indicating the loss of Lrig3 leads to an expansion of the colonic stem cell compartment, in an Erk-dependent manner. Our study describes the expression of Lrig3 within the colon, defines perturbations in mice lacking *Lrig3*, and supports a role for Lrig3 in the establishment of both colonic crypt structure and cellular census, defined as the epithelial cell type and number in colon crypts.

**Graphical Abstract:** 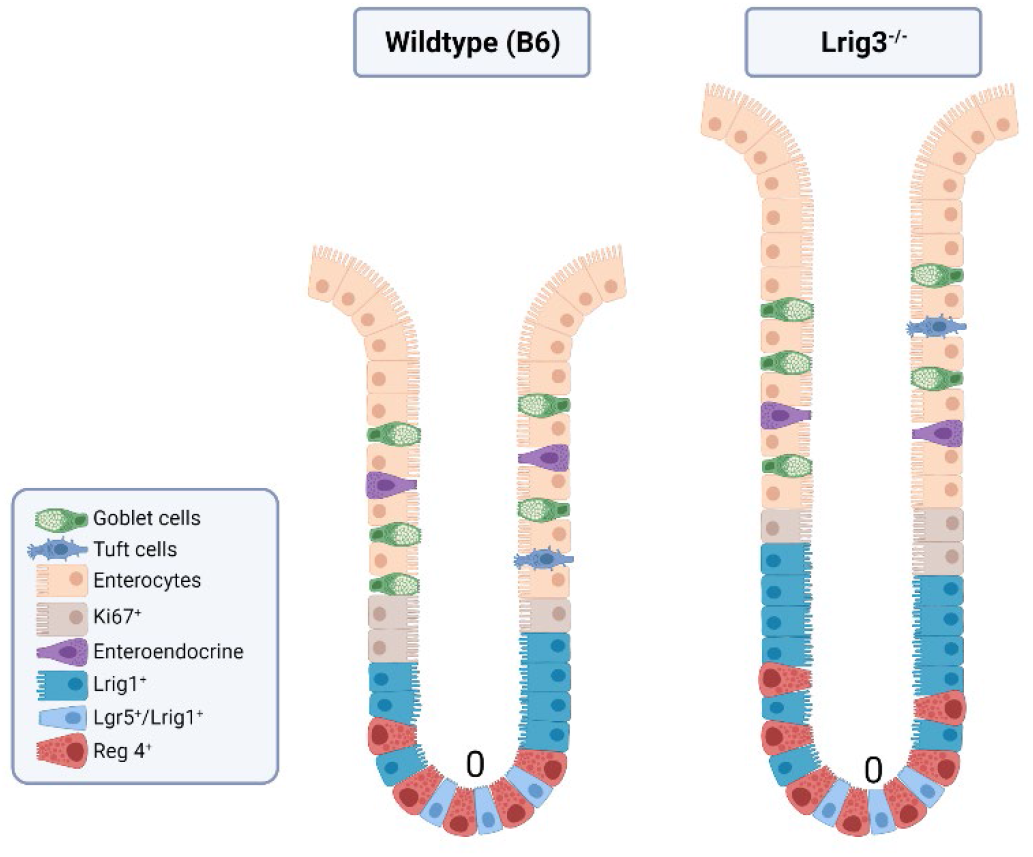

## Introduction

Precise spatiotemporal regulation of stem cell proliferation, differentiation, and apoptosis are critical for proper regeneration of the gastrointestinal epithelium. In the mouse colonic epithelium, the majority of cells are renewed every five days driven by stem cell division at the base of the colonic crypt (10). In order to maintain crypt size and epithelial cell number, the cellular census is tightly regulated by balancing the number of stem, progenitor, and differentiated cells per crypt. We know that each of these cell types are key for the absorptive and secretory jobs of the colon (13). Here, we evaluate how the loss of Leucine-rich repeats and immunoglobulin-like domains 3 (*Lrig3*) affects crypt cellular census and explore the impact of the loss of Lrig3 on these three critical cell types.

Lrig3 is a single-pass transmembrane protein and one of three members of the Lrig protein family (22). Lrig family members generally regulate both receptor tyrosine and serine/threonine kinases (22). To date, Lrig3 hasn’t been as extensively studied as the other Lrig family members, yet we know it is expressed across endo-, meso- and ectodermal tissue types throughout embryonic development (8). Disruptions in Lrig3 affect proper neural crest formation in *Xenopus laevis* and result in incomplete formation of the murine lateral semicircular canal (2, 30). Generally, mice, sheep and pig are smaller if they have a loss-of-function in Lrig3 (1, 11, 15). Only recently have contributions to primary literature uncovered mechanistic roles for Lrig3 in the progression of many cancer cell types: cervical, hepatocellular, prostate, glioblastoma and colon (5, 6, 12, 17, 23, 24, 28).

While most adult vertebrates can survive and maintain some sort of homeostasis with the loss of Lrig3, there are a number of defects that have been described in knockout mice, including cardiac hypertrophy, misregulation of insulin levels and the misorganization of neurons in the central nervous system (7, 11). In certain tissues, Lrig3 works in concert with another family member, Lrig1 (7, 8). In the gastrointestinal tract, Lrig1 is expressed on stem and progenitor cells in the small intestine and colon and is functionally important for maintaining homeostasis. The loss of Lrig1 results in aberrant growth factor signaling and tumor formation (18, 25). From publicly-available expression databases, we know Lrig3 is also highly expressed in the gastrointestinal tract (32) and Lrig3 protein is predicted to have striking homology to Lrig1 (22). These reasons led us to hypothesize that Lrig3 is important for development and homeostasis of the mouse colon. The goal of our study was to understand the impact of the loss of Lrig3 on colon biology. Through transcript and protein expression analysis, we show that Lrig3 is present throughout the colonic crypt epithelium, and we describe multiple colon morphological and cellular defects in *Lrig3*^*-/-*^ mice. Our study describes the expression of Lrig3 within the colon, defines perturbations in mice lacking *Lrig3*, and supports a role for Lrig3 in the establishment of both colonic crypt structure and cellular census, defined as the epithelial cell type and number in colon crypts.

## Results

### Lrig3 expression in colonic crypts

Our first step was to define Lrig3 expression in the colonic crypts by assessing *Lrig3* transcript and Lrig3 protein expression via *in situ* hybridization, immunofluorescence, and western blot analysis, respectively. We generated an *Lrig3* knockout mouse (*Lrig3*^*-/-*^, Fig. 1*A* and Supplemental Fig. S1) to both serve as a control for these analyses and to evaluate the impact of the loss of *Lrig3*. Comparing wildtype (WT) tissue to *Lrig3*^*-/-*^ tissue, we observed *Lrig3* transcript expression in all colonic epithelial cells and this expression was lost in the *Lrig3*^*-/-*^ mice (Fig. 1*B-E*). Further, Lrig3 protein is expressed in the majority of epithelial cells; this was confirmed by immunofluorescence and western blot analyses. Lrig3 protein was absent in *Lrig3*^*-/-*^ mice (Fig. 1*F-J*). These results define, for the first time, Lrig3 expression in the mouse colonic crypt.

**Figure 1.**
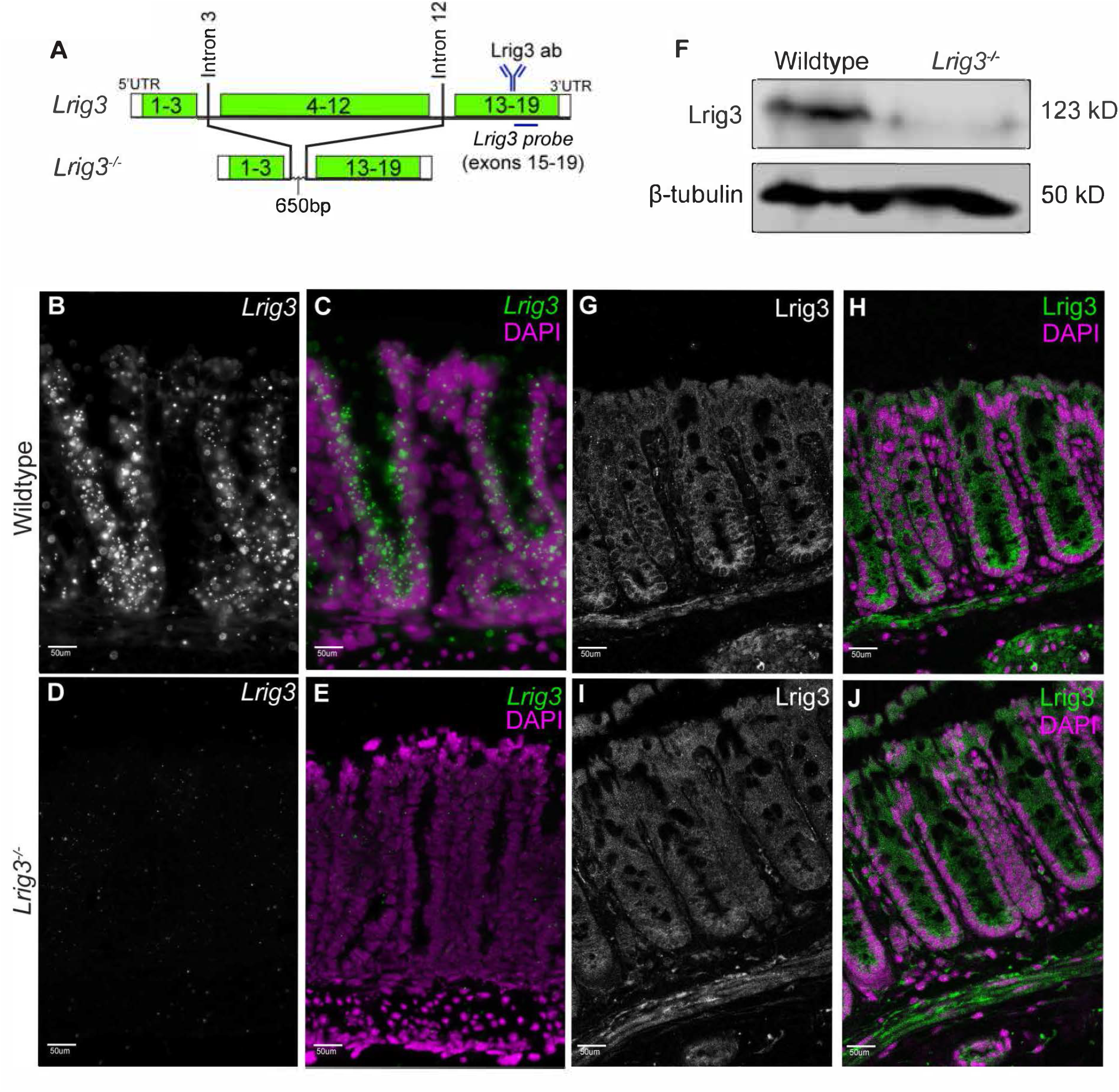
Lrig3 expression in the colonic epithelium. A. Schematic depicting knockout of exons 4-12 in the *Lrig3*^*-/-*^ mice, compared to WT mice. Blue antibody drawing indicates where Lrig3 antibody (panels F, G-J) antigen is located. Blue line indicates where *Lrig3* RNA SCOPE probe (panels B-E) is located. B-C. Representative epifluorescence RNA SCOPE images of colonic tissue cross sections indicating *Lrig3* transcripts (green) are located largely within the colonic epithelial compartment in wildtype mice. In *Lrig3*^*-/-*^ mice (D-E), RNA SCOPE signal (green) is lost (n=3 mice/genotype, 16 images/mouse). F. Western blot comparing Lrig3 antibody reactivity in wildtype and *Lrig3*^*-/-*^ colonic epithelial cell isolation lysates (n=3). G-J. Representative images from a single confocal slice illustrating Lrig3 antibody reactivity with wildtype (G-H) colonic tissue (green) cross-sections (n=3 mice/genotype, 10 images/mouse). Background fluorescence observed in Lrig3^-/-^ (I-J). Scale bars indicate 50um. Nuclei in B-E and G-J are depicted in magenta. B, D, G and H are single channel representations of the green color shown in C, E, H and J, respectively.

### *Lrig3*^*-/-*^ mice have morphological and epithelial defects

As our initial results demonstrate that Lrig3 is highly expressed throughout the colonic crypt, we posited there may be epithelial disruption to the crypts in mice lacking *Lrig3*. Indeed, initial survey of the colonic tissues indicated morphological differences, as well as epithelial cellular census differences between WT and *Lrig3*^*-/-*^ mice. To examine this directly, we first assessed colon lengths, and found that adult *Lrig3*^*-/-*^ mice have significantly longer colons than WT mice at 6-10 weeks of age (p<0.05, Fig. 2*A*). We hypothesized that along with the observed increase in colon length there may be alterations at the cellular level. We examined this by measuring total mucosal area of WT and *Lrig3*^*-/-*^ mice using hematoxylin and eosin stained, paraffin-embedded tissue sections and found that *Lrig3*^*-/-*^ mice have a significantly larger mucosal area (p<0.01, Fig. 2*B-D*), per tissue section. The larger mucosal area in *Lrig3*^*-/-*^ mice could be for several reasons, but we prioritized the investigation of two obvious hypotheses: 1) the epithelial cells within *Lrig3*^*-/-*^ mice are larger or 2) there are more cells present in the crypts of *Lrig3*^*-/-*^ mice. After an extensive examination of the colon sections from both sets of mice, we determined there was no obvious difference in cellular size between the cohorts (data not shown), ruling out that hypothesis. To investigate our second hypothesis, we quantified the number of epithelial nuclei per crypt, across both cohorts, and found that *Lrig3*^*-/-*^ mice have more cells per colonic crypt than the WT mice (p<0.05, Fig. 2*E*). While there could be many reasons for a greater number of cells, we first examined proliferation in the crypt of *Lrig3*^*-/-*^ mice compared to WT mice by quantifying the expression of Ki67, a proliferative marker, in the epithelial cells. Surprisingly, we found no significant difference in the number of proliferating epithelial cells in the colonic crypts of *Lrig3*^*-/-*^ mice at 6-10 weeks of age, compared to WT mice of the same age (Fig. 2*F-J*). From these analyses, we conclude that *Lrig3*^*-/-*^ mice have a greater number of cells per crypt, but these cells are not undergoing aberrant proliferation.

**Figure 2.**
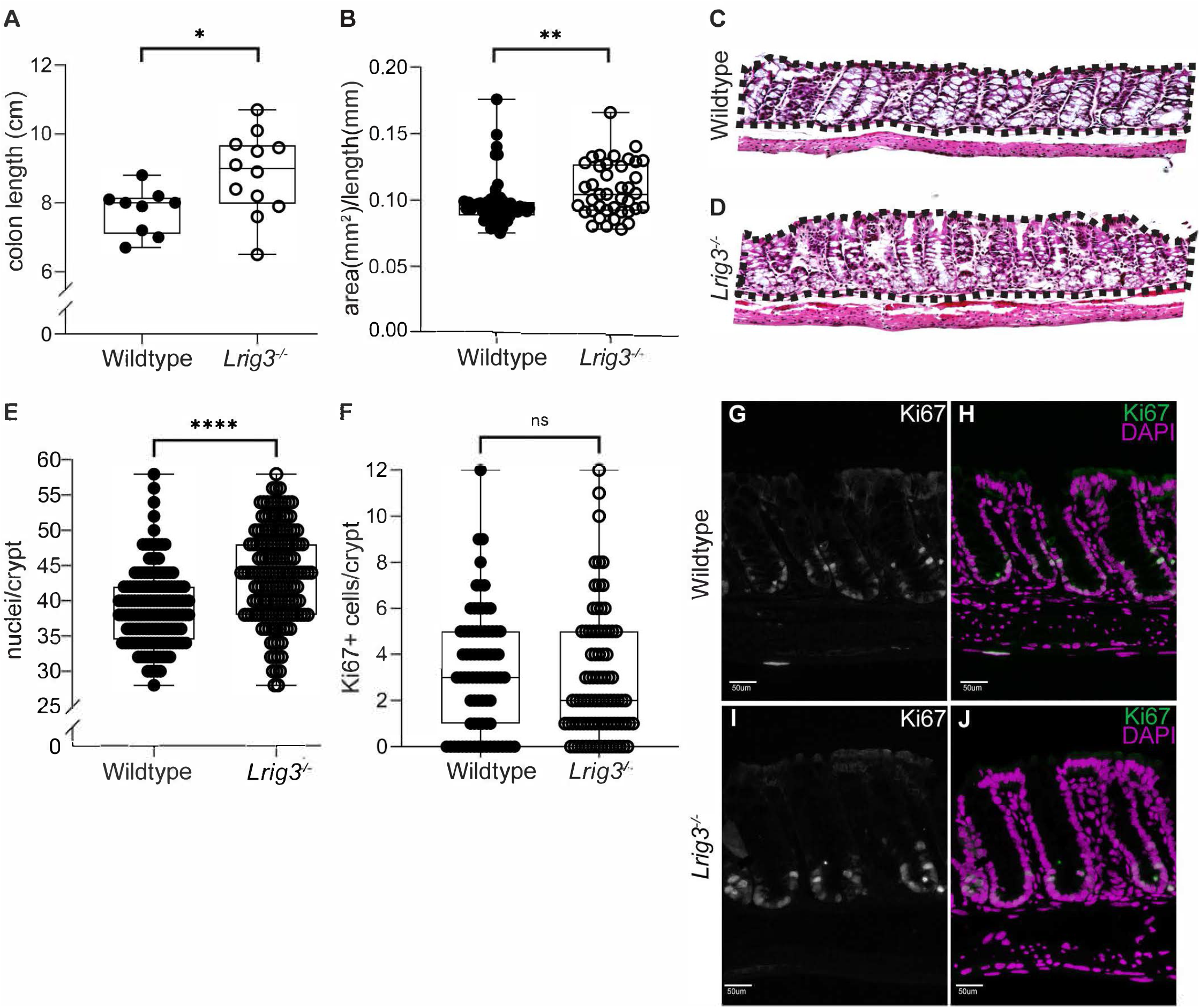
*Lrig3*^*-/-*^ mice have expanded mucosal area. A. Scatter plot indicating adult *Lrig3*^*-/-*^ mice have significantly longer colons, as measured from cecum to rectum, compared to wildtype mice (n=10 mice/genotype). B. Scatter plot indicating adult *Lrig3*^*-/-*^ mice have significantly more mucosal area in their colon, compared to wildtype mice (n=4, 10 images/mouse). C-D. Representative colon cross sections of wildtype and *Lrig3*^*-/-*^ mice that have been stained with hematoxylin and eosin. Dashed line indicates the area measured for mucosal area. E. Scatter plot indicating adult *Lrig3*^*-/-*^ mice have more nuclei per crypt compared to WT mice (n=4 mice/genotype, 10 images/mouse). F. Scatter plot indicating no significant change in proliferation (Ki67) between *Lrig3*^*-/-*^ and wildtype mice (n=4 mice/genotype, 10 images/mouse, 152 crypts/genotype). G-J. Representative images of colonic tissue sections with anti-Ki67 antibody reactivity in *Lrig3*^*-/-*^ (I-J) and wildtype (G-H) adult animals. Nuclei in H and J are depicted in magenta. In all plots, error bars indicate standard deviation from the mean. Significance was determined using an unpaired t-test where a significant difference between the groups is represented by an asterisk (*) when p<0.05. When p<.01., the data have been labeled with two asterisks (**), and by (****) when p<0.0001. Scale bar in C-D indicates 50um. All breaks in the Y axes are shown with parallel lines. G and I are single channel representations of the green color shown in H and J, respectively.

### Colon crypt stem compartments are expanded in *Lrig3*^*-/-*^ mice

As epithelial cell number expansion is a hallmark of *Lrig3*^*-/-*^ crypts, we wondered if this increased cell number was attributed to a particular region of the crypt. As there are many cell types present, we first assessed the stem and progenitor cells in the crypt-base by examining *Lgr5* expression, as this is indicative of the presence of long-lived proliferative stem cells in the base of the colonic crypt (4). First, we performed RNA Scope *in situ* hybridization for *Lgr5* expression and identified significantly more *Lgr5+* cells in the colonic crypts of *Lrig3*^*-/-*^ mice (Fig. 3*A-D*). In addition, some cells seemed to harbor qualitatively more *Lgr5* than others, even in the WT tissue, consistent with previous analyses of stem and progenitor markers in the gut (10). To more carefully examine this observation, we devised a quantification scheme for looking at *Lgr5* transcript expression levels in each cell. We binned them according to low- (<6 puncta per cell), mid- (6-14 puncta per cell), and high (15≤ puncta per cell) ranges (Fig. 3*E*). We quantified the puncta in *Lrig3*^*-/-*^ and WT cells and while we observed no difference in the number of *Lgr5* low-expressing cells, there were significantly more mid- and high-expressing cells in *Lrig3*^*-/-*^ mice (p<0.001, Fig. 3*F*). The other cell type present in the stem cell region of the crypt is defined by the expression of the Reg4 protein. As Reg4+ cells act as support cells for Lgr5+ cells (20), we reasoned that we may also observe a parallel increase in Reg4+ cells, in *Lrig3*^*-/-*^ mice. To determine this, we quantified the number of Reg4+ cells and found significantly more Reg4+ cells in the base of the crypt (p<0.0001, Fig. 3*G-K*). Reg4 is also expressed by secretory cells near the top of the crypt, thus we quantified the number of Reg4+ cells in the upper half of the crypt and determined there is no change in the number of these cells between *Lrig3*^*-/-*^ and WT mice (Fig. 3*L*). Finally, we examined the number of Vil-1+ (absorptive), Dclk1+ (Tuft), and ChgA+ (enteroendocrine) cells, which correspond to three additional differentiated cell types found in the upper portion of the crypts. Qualitatively, we found no obvious difference in Vil-1 expression, when comparing approximately 50 images per genotype. Quantitatively, we found no difference in Dclk1+ and ChgA+ cells when comparing *Lrig3*^*-/-*^ and WT mice (Supplemental Fig. S2). Collectively, our results demonstrate the increased cell number we observe in *Lrig3*^*-/-*^ mice is restricted to stem and support cells in the crypt base.

**Figure 3.**
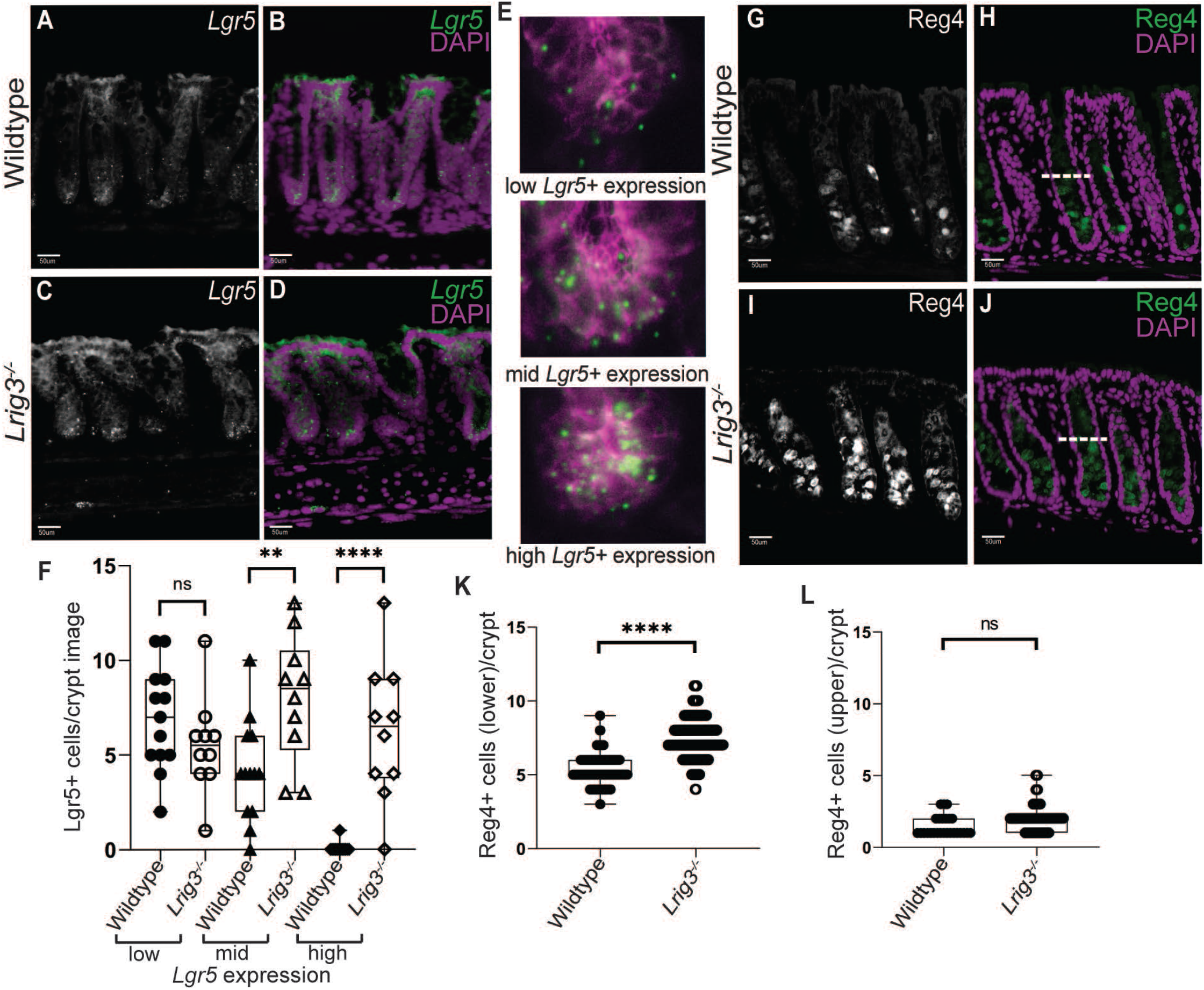
Expansion of the stem cell niche. A-D. Representative RNA SCOPE images of colonic tissue cross sections indicating *Lgr5* transcripts (green) between wildtype (A-B) and *Lrig3*^*-/-*^ (C-D) mice. E. Representative images of low, mid, and high *Lgr5+* transcript expression. F. Scatter plot indicating *Lrig3*^*-/-*^ mice have significantly more *Lgr5*^*+*^ mid and high cell transcript expression compared to wildtype mice (n=3 mice/genotype, 4 images/mouse). G-J. Representative images of colonic tissue sections comparing expression of Reg4 antibody reactivity. Dotted line depicts the center of the crypt where they were marked and quantified crypt upper half and lower half. K. Scatter plot indicating an increase in Reg4+ cells, in the lower half of the colonic crypt, in *Lrig3*^*-/-*^ mice compared to wildtype mice (n=7 mice/genotype, 7 images/mouse). L. Scatter plot indicating no change in Reg4+ cells in the upper portion of the colonic crypt of Lrig3^-/-^ mice compared to wildtype mice (n=7 mice/genotype, 7 images/mouse). Nuclei in B-E are depicted in magenta. Scale bar indicates 50um. In all plots, error bars indicate standard deviation from the mean. Significance was determined using an unpaired t-test where a significant difference between the groups is represented by an asterisk (*) when p<0.05. When p<.01., the data have been labeled with two asterisks (**), when p<0.0001, the data have been labeled with four asterisks (****). A, C, G and I are single channel representations of the green color shown in B, D, H and J, respectively.

### *Lrig3*^*-/-*^ mice have increased Lrig1 and decreased in pErk

As the stem cell niche is expanded in *Lrig3*^*-/-*^ mice, we next examined if there was any change in expression of Lrig1, which is an Lrig3 family member, and widely expressed in intestinal stem and progenitor cells(18). While Lrig1 is normally restricted to the crypt-base, we found that *Lrig3*^*-/-*^ mice have more Lrig1+ cells in the mid- and upper-crypt (Fig. 4*A-D*). We quantified the number of Lrig1+ cells per crypt and found significantly more Lrig1+ cells in *Lrig3*^*-/-*^ mice compared to WT mice (p<0.0001, Fig. 4*E*). We also quantified Lrig1 expression in the crypts via western blot analysis and confirmed the increase in Lrig1 we detected in *Lrig3*^*-/-*^ mice (Fig. 4*F*). As loss of Lrig1 results in changes to growth factor receptor signaling (18, 26), we next assessed if the over-expression of Lrig1 seen in *Lrig3*^*-/-*^ mice resulted in cell signaling changes in the crypts. We prioritized analysis of total and activated Epidermal growth factor receptor (Egfr and pEgfr, respectively), as these receptors are modulated based on Lrig1 expression (18). We also examined extracellular signal-related kinase (Erk) expression, as this is a transcriptional activator of proliferation and differentiation, and promoter of stemness in both the small intestine and colon (16). We observed no obvious differences in the protein expression of total Egfr (Fig. 4*G-H, K*), activated Egfr or total Erk (Fig. 4*I-K*); however, we consistently observed decreased expression of phosphorylated Erk (pErk) in the *Lrig3*^*-/-*^ colonic crypts by western blot analysis (Fig. 4*K*). These results indicate this common downstream mediator of mitogen activated kinase signaling is changed in mice lacking *Lrig3*. Taken together, we observe that *Lrig3* is required for proper Lrig1 expression and appropriate Erk activation.

**Figure 4.**
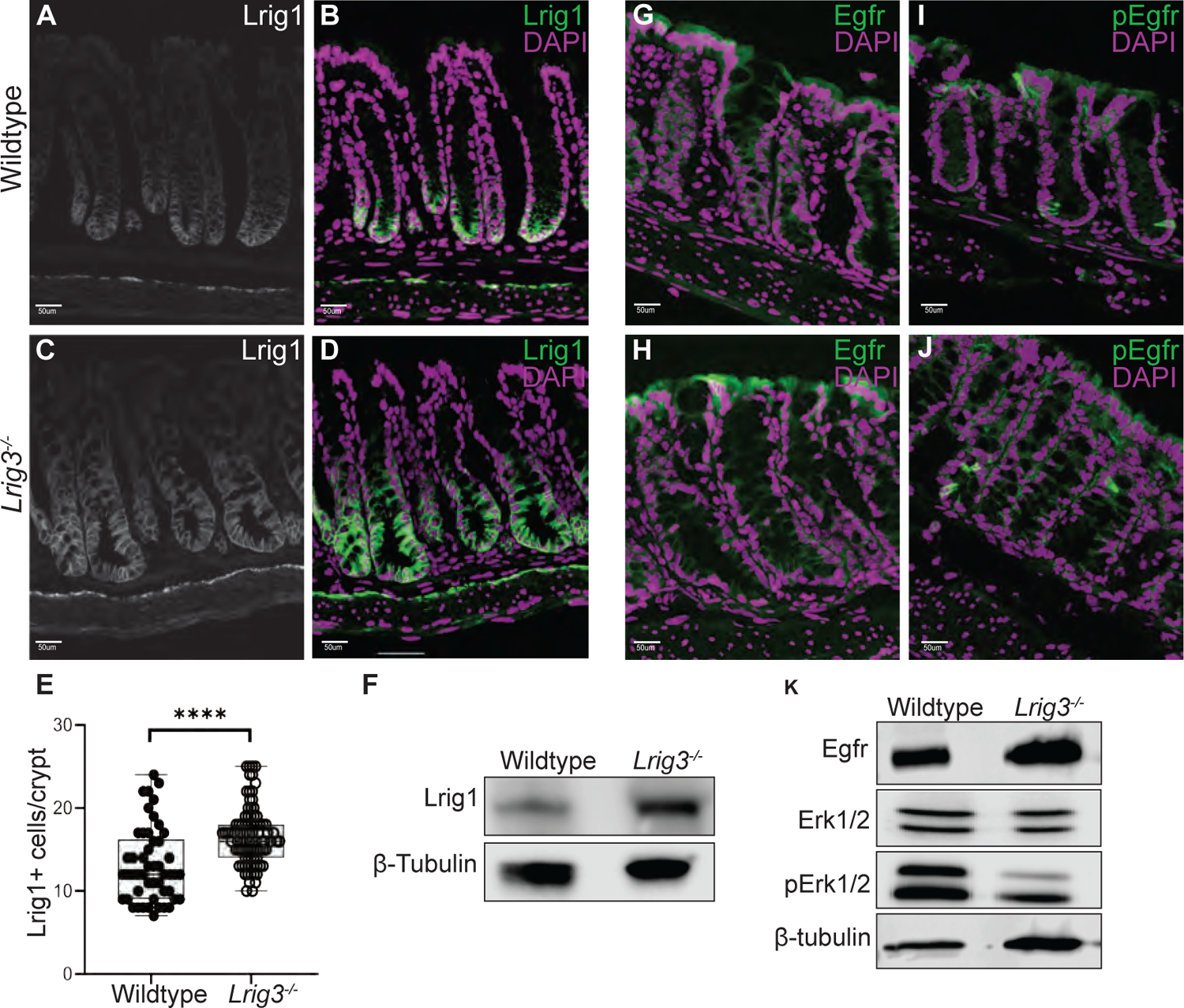
Expression of Lrig1 is expanded in *Lrig3*^*-/-*^ mice. A-D. Representative images of colonic tissue sections comparing the expression of Lrig1 between wildtype (A-B) and *Lrig3*^*-/-*^ (C-D) mice. In A and C, Lrig1 (white) is detected at the cell membrane of cells located in the crypt base. In B and D, Lrig1 (green) is shown in the context of all cells in the crypt (magenta). E. Scatter plot indicating significantly more Lrig1^+^ cells in the colonic crypt of Lrig3^-/-^ mice compared to wildtype (n=4 mice/genotype, 10 images/mouse). F. Western blot comparing Lrig1 antibody reactivity in wildtype and *Lrig3*^*-/-*^ colonic tissue cell lysates (n=3). G-J. Representative images of colonic tissue sections comparing the expression of pEgfr (I and J, green) and Egfr (G-H, green) between wildtype and *Lrig3*^*-/-*^ mice. K. Representative western blots comparing protein expression for Egfr, Erk1/2, and pErk1/2 in wildtype and *Lrig3*^*-/-*^ colonic tissue cell lysates (n=3). Nuclei in G-J are depicted in magenta. Scale bars indicate 50um. Significance in scatter plot was determined using an unpaired t-test where a significant difference between the groups is represented by an asterisk (****) when p<0.0001. A and C are single channel representations of the green color shown in B and D respectively.

## Discussion

Using our novel *Lrig3*^*-/-*^ mouse, we demonstrate that *Lrig3*^*-/-*^ mice have more cells per crypt, a greater mucosal area, and longer colons compared to wildtype mice. This suggests the expression of Lrig3 is required for both the total number of epithelial cells in the mouse colon, as well as establishment of colon length. Further, we show *Lrig3*^*-/-*^ mice have an Erk-associated expansion of the colonic stem cell compartment, as they harbor significantly more stem, progenitor, and deep crypt secretory cells.

Our research defines the expression domain of Lrig3, in the colon for the first time, as a novel marker of all colonic epithelial cells. The most well-studied Lrig family member, Lrig1 (18), is a homologue of Lrig3 with redundant expression patterns in some tissues (7, 8). In the colon, Lrig1 expression is distinct from that of Lrig3, as it is restricted to the crypt-base (18, 26). As their expression patterns are distinct in the gastrointestinal tract, it is likely that Lrig1 and Lrig3 also have unique molecular functions. *In vitro*, we know that Lrig3 stabilizes the ErbB receptors, whereas Lrig1 promotes ErbB receptor degradation (18). It is less clear if Lrig3 also promotes stabilization of ErbB receptors *in vivo*; this observation may be tissue-dependent (3). As with Lrig1, Lrig3 has been evaluated across many tissue types; and has a significant impact on growth factor receptor pathways. We show that one common mediator of ErbB and Ras signaling pathways, Erk, is dysregulated in *Lrig3*^*-/-*^ mice. Through our examination of the ErbB family member, Egfr, we did not observe any changes to expression or activation of this powerful growth factor receptor in *Lrig3*^*-/-*^ mice; the impact of Lrig3 loss on other growth cell signaling pathways represents an important area of future research. Lrig3 also has been shown to regulate fibroblast growth factor (FGF) receptors which results in changes to neural crest formation in frog (30) and vascular endothelial growth factor (VEGF) receptors are modulated by Lrig3 in glioma (17, 31). Both FGF and VEGF pathways are important for the development and maintenance of the colonic epithelium (9, 21), but it is unknown if Lrig3 modulates these pathways in this context. This represents an important area of future research.

The question of how Lrig3 is regulated may be equally as important as the question as to what Lrig3 is regulating. Notably, *Lrig3* expression is directly modulated by miRNA-196a, a microRNA that is upregulated in gastric, colon, breast and pancreatic cancer and promotes tumorigenesis. In cervical cancer cells, miRNA-196a targets Lrig3 (19) and improves cell viability. Lrig3 is also regulated at the post-transcriptional level, as there are reports of a circular form of Lrig3 *(circRNA Lrig3*) that is highly expressed in human hepatocellular carcinoma (HCC). Detection of blood plasma *circ-Lrig3* is a highly sensitive and specific diagnostic indicator for HCC (23). Mechanistically, downregulation of *circRNA Lrig3* represses both the MAPK/ERK and Smad pathways to prevent the progression of HCC and the loss of *circRNA Lrig3* suppresses proliferation, invasion of HCC cells *in vitro* and blocks tumor growth of HCC *in vivo* (12, 24). One recent report gives us a glimpse of the potential role in advanced colon cancer. Using human colon cancer cell lines, Zang and colleagues were able to show that Lrig3 represses metastasis-associated cell motility by inhibiting slug via inactivating ERK signaling (29). While the data on Lrig3 expression in solid tumors is expanding rapidly (5, 6, 12, 17, 23, 24, 29), the role of Lrig3 in the gastrointestinal tract at the genome, transcriptional and protein level is largely unknown and represents a valuable avenue for future research for both normal and cancerous colon tissue.

In terms of the requirement for Lrig3 in colon development and homeostasis, we have two key observations. The first is that *Lrig3*^*-/-*^ mice are born physically smaller and weigh less than their WT counterparts (11), yet we also consistently observe these mice to have significantly longer colons at adulthood. While there has been a great deal of research defining the molecular requirements for small intestinal morphogenesis, and to some extent, for the colon (9, 13), our results clearly signify the importance of studying Lrig3 and its role in colon development. While it is beyond the scope of this paper, a clear next step to understanding how *Lrig3* impacts colon development is to take an *in vivo*, inducible loss-of-function approach for Lrig3, using a timecourse strategy to define the impact of loss of Lrig3 on birth weight and colon length.

Perhaps the most striking observation from our examination of *Lrig3*^*-/-*^ mice is the greater number of cells per crypt despite no significant change to proliferation, resulting in *Lrig3*^*-/-*^ mice having an expanded pool of stem and support cells in the crypt-base. While colonic epithelial self-renewal has been extensively researched (10) the majority of this research has been in adult mice, and generally has been assessed during regenerative phases after tissue injury (27). From these studies, we know the key signaling pathways that promote stem cell self-renewal and daughter cell differentiation (10). Cellular census (epithelial cell type and number) is generally consistent from crypt-to-crypt in the distal colon of both mice and humans. Unfortunately, there is a paucity of research examining the molecular factors that regulate the number of colon cells and how the crypts arrive at a homeostatic census (and thus a consistent crypt size) in adulthood. Our data clearly demonstrate the requirement for proper Lrig3 expression to yield appropriate crypt cellular census. Despite the increase in cell number per crypt in *Lrig3*^*-/-*^ mice, we do not observe an increase in all epithelial cell types. Indeed, we observe a specific increase in the number of cells in the stem cell compartment at the crypt-base, suggesting that *Lrig3* is critical for establishing the size of the stem cell compartment during development. In future experiments, it will be important to test this hypothesis directly, either by overexpression of Lrig3 or through forced, wide-spread regeneration of colon crypts in *Lrig3*^*-/-*^ mice. In the latter example, induction of acute injury to the adult *Lrig3*^*-/-*^ colon to spur crypt renewal will be instructive in understanding how *Lrig3*^*-/-*^ mice arrive at this aberrant state of homeostasis. Taken together, our results signify the importance of studying Lrig3 and its role in cellular census and colon development.

## Experimental Procedures

### Generating *Lrig3*^*-/-*^ mice

The *129 Lrig3* BAC clone (bMQ-129G9) was obtained from Sanger Institute. The *Lrig3* targeting construct which disrupts exons 4-12 of the Lrig3 locus was generated by BAC recombineering (14). The gene disruption strategy, null allele sequence and location of PCR primers is presented in Supplemental Figure 1. PCR primer sequences are listed in Supplemental Table 1. ES cell electroporation and subsequent blastocyst injections were performed by the Transgenic Mouse/ES Cell Shared Resource at Vanderbilt University.

### Mice

C57BL/6 and *Lrig3*^*-/-*^ mice were housed in a specific pathogen-free environment under controlled light cycle conditions, fed standard rodent lab chow, and provided water *ad libitum*. All procedures were approved and performed in accordance with the University of Oregon Institutional Animal Care and Use Committee. All mice were used at 6-10 weeks and mouse sex was mixed male to female at a roughly 50% ratio of each. All mice were sacrificed by cervical dislocation. At time of sacrifice, colons were removed, flushed with ice cold PBS and immediately measured using a ruler to obtain colon length, and then bifurcated.

### Tissue Preparation for Staining

Tissue for paraffin and frozen block preparation were pinned onto a wax surface, fixed using 4% paraformaldehyde (PFA) for one hour (tissue imaged in Fig1*G-J* were fixed in 4% PFA overnight), on a shaker at room temperature. They were then washed three times (five minutes each) in PBS. For frozen blocks, tissue was submerged 30% sucrose in PBS overnight at 4°C and embedded in optimal tissue compound (OCT) for subsequent sectioning. For paraffin blocks, tissue was incubated in 70% ethanol and dehydrated in increasing alcohol baths and embedded in paraffin wax. All slides were sectioned at 7µm, (unless stated otherwise), and stained according to procedures below.

### Colonic Crypt Protein Isolation

Bifurcated colons were cut into ∼ 1cm pieces and incubated in ice cold 2mM EDTA and 0.5mM DTT PBS buffer, washed in PBS, and incubated 2mM EDTA buffer at 37°C. Tissue underwent vigorous lateral shaking to release crypts from submucosa, suspension was decanted, and shaking was repeated three times. The presence of single crypts is verified under a compound microscope and residual submucosal tissue is removed. The final collection of crypt suspensions was centrifuged at 500xg, cell pellet was then resuspended in Pierce RIPA buffer (ThermoFisher) with protease (½ tablet, Pierce Protease Inhibitor Mini tablets, EDTA-free, ThermoFisher) and phosphatase inhibitor (½ tablet PhosSTOP EASYpack, Roche), homogenized using an 18-gauge needle, and centrifuged at maximum speed. The supernatant was removed, and protein concentration determined using Pierce BCA kit (ThermoFisher).

### Morphometric Analysis

Mucosal area and nuclei per crypt were quantified using paraffin embedded tissue and stained with Hematoxylin and Eosin (VWR). Images and measurements were obtained on a Nikon Eclipse/Ds-Ri2. To quantify the mucosal area we measured 10 images per animal (n=4 mice per genotype) using the Nikon NIS-Elements measurement tool. We measured the area of colonic epithelium by drawing lines across the basement membrane of the epithelium, along the sides, and across the luminal edge. To quantify the nuclei per crypt we used Hematoxylin and Eosin-stained paraffin slides and counted total nuclei per crypt in 10 images per animal (n=4 mice per genotype).

### Antibodies and Staining Procedure

Frozen tissue slides were washed in PBS three times (three minutes each), blocked in 1% bovine serum albumin (BSA) and 0.03% Triton X-100 suspended in PBS for 1 hour. Antibodies were diluted in this blocking buffer at concentrations listed in Supplemental Table 1, applied to the sections and sections were incubated overnight at 4°C. Slides were then washed in PBS three times (three minute each) and incubated with secondary antibodies at 1:500 in the same blocking buffer as above, for one hour at room temperature. Lastly, slides were washed in PBS for three minutes, then washed in PBS plus DAPI (1:10,000) for five minutes, and finally washed in PBS for three minutes. Slides were mounted using an n-propyl gallate/glycerol solution. Paraffin slides underwent conventional deparaffinization in xylenes and rehydration via ethanol washes and water, followed by antigen retrieval in a 1x Citrate buffer (ThermoFisher) in a pressure cooker for one hour. Antibody catalog information and concentrations used are listed in Supplemental Table 2.

### Western Blot

Protein was diluted in a Laemmli buffer with 5% β-mercaptoethanol and boiled at 95°C for five minutes. 25µg of protein was loaded per lane into an SDS-PAGE gel (Bio-Rad Mini-Protean pre-cast gels 4-20%) and gels were run at 160V for approximately 90 minutes. Protein was transferred from the gel to a PVDF membrane overnight at 55V at 4°C. Membranes were blocked in either 5% dry milk or 5% BSA in PBS for one hour and incubated in primary antibody overnight at 4°C. Membranes are then incubated in HRP-conjugated secondary antibodies at room temperature for one hour. ECL reagent (GE Healthcare) was then applied and subsequent images were captured on a LI-COR Odyssey Fc Imaging System. For quantification of protein levels of target proteins, β-tubulin was used as a normalization control (n=3 mice per genotype).

### *In Situ* hybridization

*In situ* hybridization was conducted using RNAscope Multiplex Fluorescent Reagent Kit v2 (Advanced Cell Diagnostics #323110) according to manufacturer protocols for fixed, frozen tissue sample preparation protocol (ACD TN 320535 Rev A, 323100-USM). Briefly, 15um frozen sections were washed briefly in PBS, boiled for 5 minutes in 1x Target Retrieval buffer, followed by two brief washes in ddH_2_O and one in 95% EtOH. Slides were dried then dammed with an ImmEdge hydrophobic barrier pen, then incubated with Protease III for 15 minutes at 40°C. Slides were washed briefly in ddH_2_O, then incubated at 40°C in sequence with Probe-Mm-Lgr5 (312171) or or Mm-Lrig3 probe (310541) (2hr), AMP 1 (30 min), AMP 2 (30 min), AMP 3 (15 min), HRP-C1 (15 min), Opal-570 TSA (30 min, 1:1000), with two two-minute washes with 1x Wash Buffer between hybridization steps. Slides were counterstained with DAPI and mounted with N-propyl gallate mounting medium.

### Fluorescence microscopy: image acquisition and analysis

Images for quantification of all immunofluorescent and RNAscope images were obtained using a Nikon Eclipse/Ds-Ri2 and NIS Elements software tools. Lrig3 antibody was visualized using confocal microscopy on a Zeiss LSM-880 system. Images were false-colored in Adobe Photoshop. All image analysis and quantification was performed in a double-blind fashion and statistical comparisons were analyzed using GraphPad Prism software. Quantification of all images (total “n” and counting metrics) are designated in each figure legend, however there were two sets of images analyzed with different criteria and we explain them here. For the quantification of *Lgr5* expression, images were acquired and binned according to low- (<6 puncta per cell), mid- (6-14 puncta per cell), and high (15< puncta per cell) expression levels. Example images are shown in Fig. 3*E*. For Reg4 quantification, we separated each colonic crypt in half (upper and lower regions) and counted the positive cells in each region. Example images are shown in Fig. *H* and *J*, with the dotted white line separating upper from lower.

## Figure Legends

**Supplemental Figure 1. Wildtype and Lrig3^-/-^ allele map, sequence, and PCR screening of mice**. A-B. Schematic representation of the wildtype and *Lrig3*^*-/-*^ genes. Disruption of *Lrig3* was achieved through the removal of exons 4-12, joining the third and twelfth intron. B. The intervening remaining sequence (650bp) is shown. C. Confirmatory PCR analysis of the wildtype *Lrig3* allele, using primers that are located upstream (5’) and downstream (3’) of the fourth exon, and generate 1.2kb PCR product. *Lrig3*^*-/-*^ mice do not generate a PCR product, as they lack the 3’ site. D. Confirmatory PCR analysis of the null *Lrig3* allele, using primers that are located upstream (5’) of the fourth exon and downstream (3’) of the twelfth exon, generates a 650bp product in the null mice. Wildtype mice do not generate a PCR product under standard conditions, as that region is 10kb in length. *Lrig3* mutant heterozygous and homozygous mice are viable and fertile. PCR primer sequences are listed in Supplemental Table 1.

**Supplemental Figure 2. Differentiated Cell marker expression in *Lrig3***^***-/-***^ **and wildtype colons**. A-D. Representative images of Vil-1 expression (green) in the absorptive cells of the colonic epithelium in wildtype (A-B) *Lrig3*^*-/-*^ colons (C-D). B’ and D’. Enlarged images of Vil-1 expression shown in wildtype (B and B’) and *Lrig3*^*-/-*^ (D and D’). E-H. Representative images of Dclk1 expression (green) in tuft cells of colonic epithelium in wildtype (E-F) *Lrig3*^*-/-*^ (G-H) colons. I. Scatter plot indicating no significant change in Dclk1^+^ cells in the colonic crypts of *Lrig3*^*-/-*^ compared to wildtype mice (n=4, 10 images/mouse). J-M. representative images of ChgA expression (green) in neuroendocrine cells of the colonic epithelium in wildtype (J-K) and *Lrig3*^*-/-*^ (L-M) colons. N. Scatter plot indicating no significant change in ChgA^+^ cells in the colonic epithelium of *Lrig3*^*-/-*^ compared to wildtype mice (n=4, 10 images/mouse). For both panels, nuclei in are depicted in magenta and the scale bar indicates 50um.

**Supplemental Table 1.**
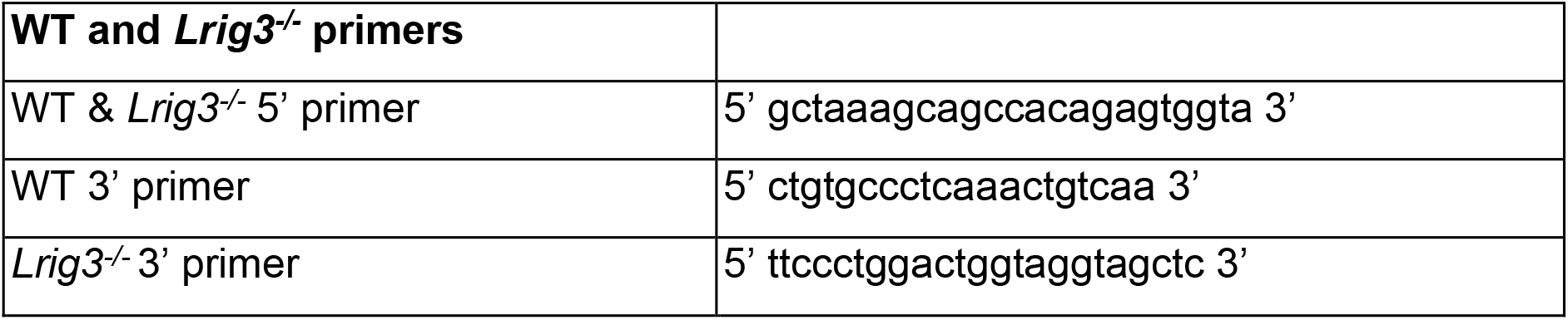

**Supplemental Table 2.**

| Antibody | Company | Product# | Concentration |
| --- | --- | --- | --- |
| Lrig3 | US Biological | 151911 | 1:250(IF)<br>1:300 (WB) |
| Lrig1 | R&D Systems | AF3688 | 1:500 (IF)<br>1:300 (WB) |
| Ki67(PE) | eBiosciences | SolA15 | 1:100 (IF) |
| Egfr | abcam | ab52894 | 1:500 (IF)<br>1:1000(WB) |
| pEgfr (Y1068) | abcam | 200709 | 1:400 (IF) |
| Erk | Bioss | BS-0022R | 1:1000 (WB) |
| pErk | Cell Signaling Technology | 4370S | 1:1000 (WB) |
| Reg4 | R&D Systems | AF1379 | 1:100 (IF) |

