## Supplementary figures and images for "Lrig3 restricts the size of the colon stem cell compartment"

### Supplemental Figures

Supplemental Figure 1

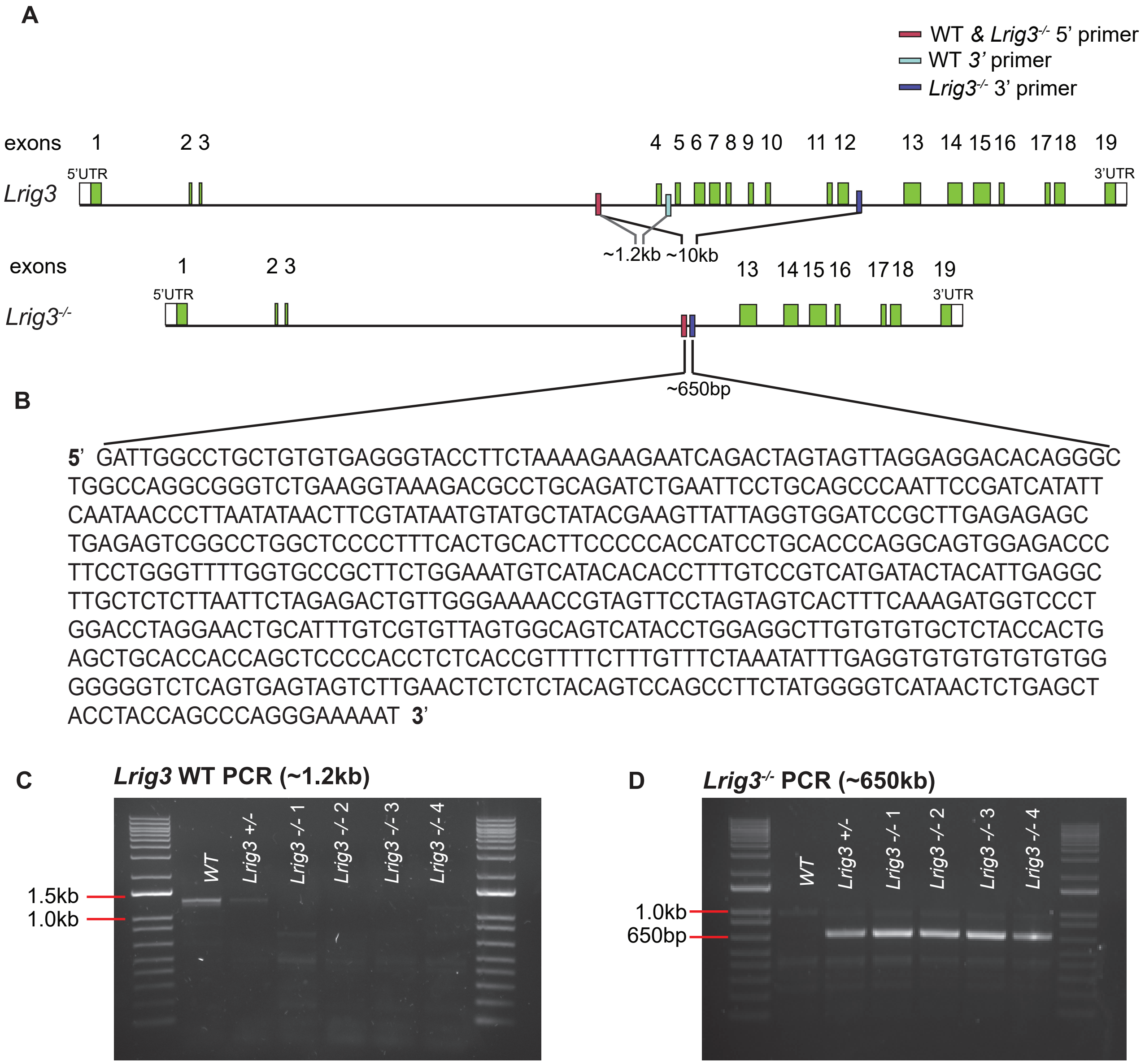

Supplemental Figure 2

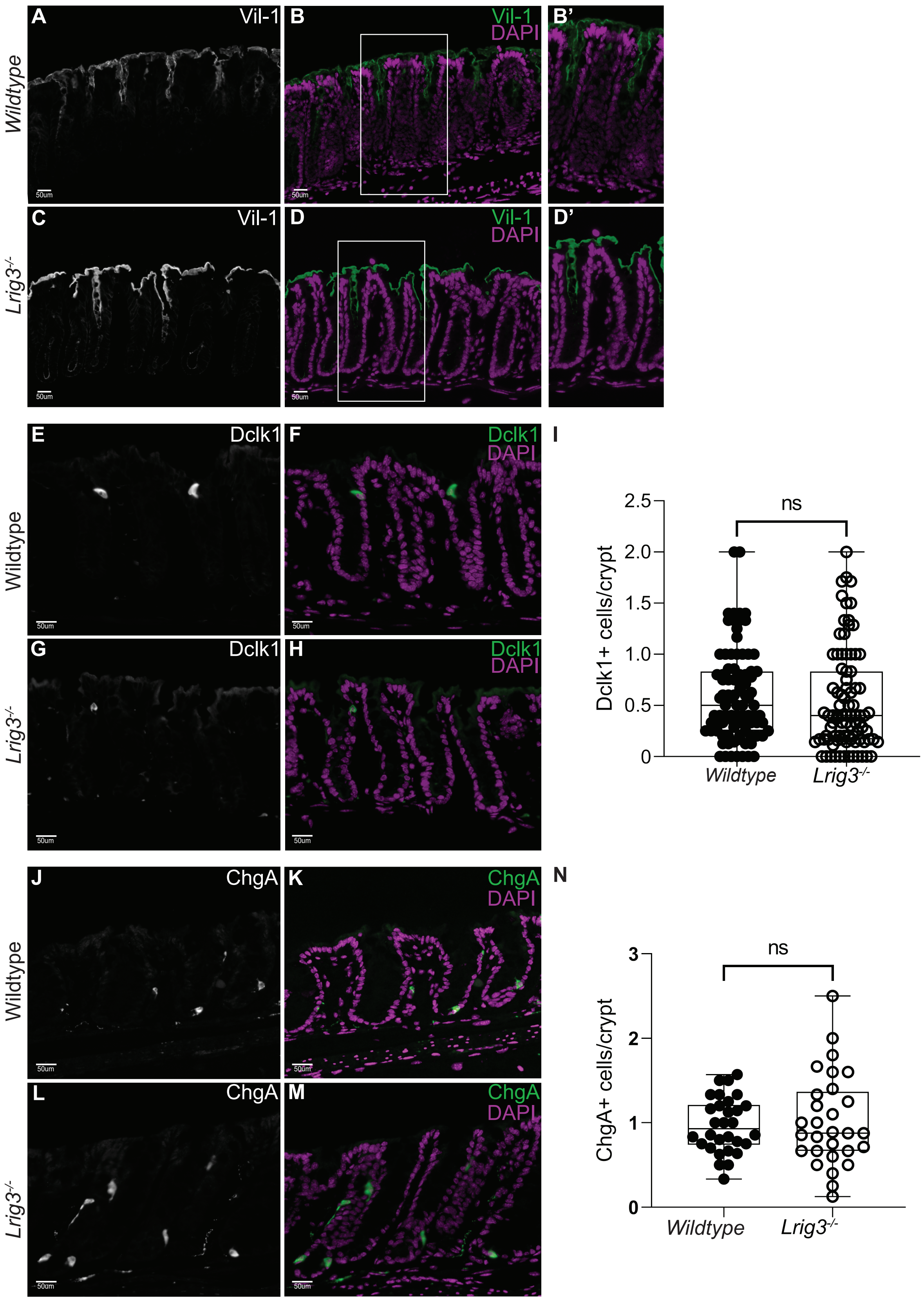
